# A holistic quantitative understanding of state transition in plant photosynthesis

**DOI:** 10.1101/2024.06.21.600050

**Authors:** Hui Ming Olivia Oung, Haniyeh Koochak, Malgorzata Krysiak, Vaclav Svoboda, Helmut Kirchhoff

## Abstract

Efficient and safe harvesting of sunlight by photosynthesis in plant thylakoid membranes requires that both photosystems (PS)I and PSII operate with similar electron turnover rates. This is realized by state transition encompassing the redistribution of light-harvesting complexes II (LHCII) between the spatially separated PSII (mainly in stacked grana thylakoids) and PSI (mainly in unstacked domains). Here, we provide a quantitative holistic view on lateral protein and pigment reorganizations within the thylakoid membrane network induced by state transitions and the role of reversible protein phosphorylation for this process. The data reveals that plants can perfectly balance electron fluxes through both photosystems by the redistribution of a certain pool of hyperphosphorylated LHCII from PSII in stacked to PSI in unstacked thylakoid membranes. Force balance analysis predicts that the photosystems antenna reorganization is not realized by a phosphorylation induced stimulation of lateral mobility of LHCII but likely by vertical unstacking. Shuffling of phospho-LHCII during state transition results in remodeling of the PSI supercomplex landscape but not of the PSII landscape supporting the notion that only a loosely bound pool of LHCIIs is involved in state transition.

## Introduction

Sessile land plants must cope with challenges dictated by an everchanging and often unpredictable environment. A particular challenge is to balance photosynthetic sunlight absorption which is realized by specialized light-harvesting complexes (LHC) in thylakoid membranes with manifold energy-demanding metabolic reactions in the plant cells. Both the energy input by solar radiation and the metabolic energy demand of the cell can fluctuate at different time scales demanding for dynamic finetuning to harmonize both processes. Failure to maintain a proper energy balance by regulatory processes has detrimental effects for plant fitness and survival in the field (Kühlheim 2002, Frenkel 2007).

One key regulatory process in thylakoid membranes is state transitions which balances the distribution of harvested sunlight between the two photosystems (Bonaventura 1969, Allen 1992). Photosystem (PS)I and PSII work in series to drive linear electron transport (LET) from water (PSII) to ferredoxin (PSI). More efficient LET is achieved when both PSs run with the same speed, i.e. their electron transport rates are synchronized. If PSII runs faster than PSI, the intersystem electron transport gets over reduced leading to photodamage. In contrast, if PSI runs faster than PSII, its extra capacity is lost leading to a waste of harvested energy from sunlight. State transition aligns the absorption cross section of the two PSs by shuffling the LHCII-bound chlorophylls (Chl) between them, synchronizing their rates.

A prerequisite for regulating the energy balance between the two PSs by LHCII redistribution is their spatial separation. This is realized by the stacking of part of the thylakoid membrane system in grana leading to stacked (mainly PSII) and unstacked (mainly PSI) domains (Anderson 1966, Staehelin 1996, Albertsson 2001) In state 1 (S1), LHCII is preferentially attached to PSII in stacked grana whereas in state 2 (S2) part of LHCII unbinds from PSII and attaches to PSI in unstacked domains to increase its antenna size. This LHCII redistribution between PSII and PSI is regulated by its reversible phosphorylation. In plants, LHCII phosphorylation is mainly catalyzed by the STN7 kinase (Bellafiore 2005, Bonardi 2005). A second kinase, STN8, mainly phosphorylates PSII phosphoproteins D1, D2, and CP43 (Bonardi 2005, Vainonen 2005) and to a minor extent LHCII subunits (Leoni 2013). LHCII dephosphorylation is catalyzed by a PPH1/TAP38 phosphatase (Pribil 2010, Shapiguzov 2010). The STN7 kinase activity is controlled by the redox state of the inter-photosystem plastoquinone pool that acts as molecular sensor for measuring imbalances in energy distribution between PSII and PSI. The STN7 kinase is inactivated at higher light intensities (Rintamäki 2000, Ancin 2019). Therefore, state transition regulates the harvesting of sunlight mainly under lower light intensities, i.e., under conditions where both PSs should run at synchronized rates for an efficient energy conversion process.

Here, we present a holistic quantitative picture on state transition demonstrating how phosphorylation-triggered lateral reshuffling of LHCII between PSII- and PSI-domains regulates antenna sizes and supercomplex formation of both photosystems and thereby their electron transport rates.

## Results and Discussion

### Different protein phosphorylation levels in stacked and unstacked thylakoid domains

The impact of state transition on protein localization, organization, and function was studied by employing a recently optimized protocol separating stacked grana and unstacked thylakoid domains (stroma lamellae + grana margins) that recovers ∼90% of proteins and Chls (Koochak 2019). At the end of the night period, plants were in S1 as indicated by the almost complete absence of LHCII phosphorylation in isolated thylakoid membranes measured by Western blot analysis with a phospho-threonine specific antibody (Fig. 1A, WT-dark). After illumination of the plants with white light of ∼120 μmol quanta m^-2^ s^-1^, LHCII phosphorylation clearly increases (Fig. 1A, WT-Light) indicative for transition to S2 (further confirmed by low temperature Chl fluorescence spectra in Fig. 4). To explore the full capacity of state transition, the kinase double mutant *stn7/8* (locked in S1, Bellafiore 2005, Bonardi 2005, Leoni 2013) and the phosphatase double mutant *pph1/pbcp* (locked in S2, Pribil 2010, Shapiguzov 2010, Samol 2012) were employed. As expected, the *stn7/8* mutant shows no protein phosphorylation both in dark and light (Fig. 1A). In contrast, the *pph1/pbcp* mutant shows significant LHCII phosphorylation in the dark due to inefficient dephosphorylation that is further increased by illumination (Fig. 1A). Since dephosphorylation is lacking in *pph1/pbcp* and the STN7 kinase is activated in the light, it is reasonable to assume that LHCII is maximally phosphorylated in *pph1/pbcp*-light. It follows that the one-hour light treatment leads to 87% LHCII phosphorylation in WT (Fig. 1A).

**Fig. 1.**
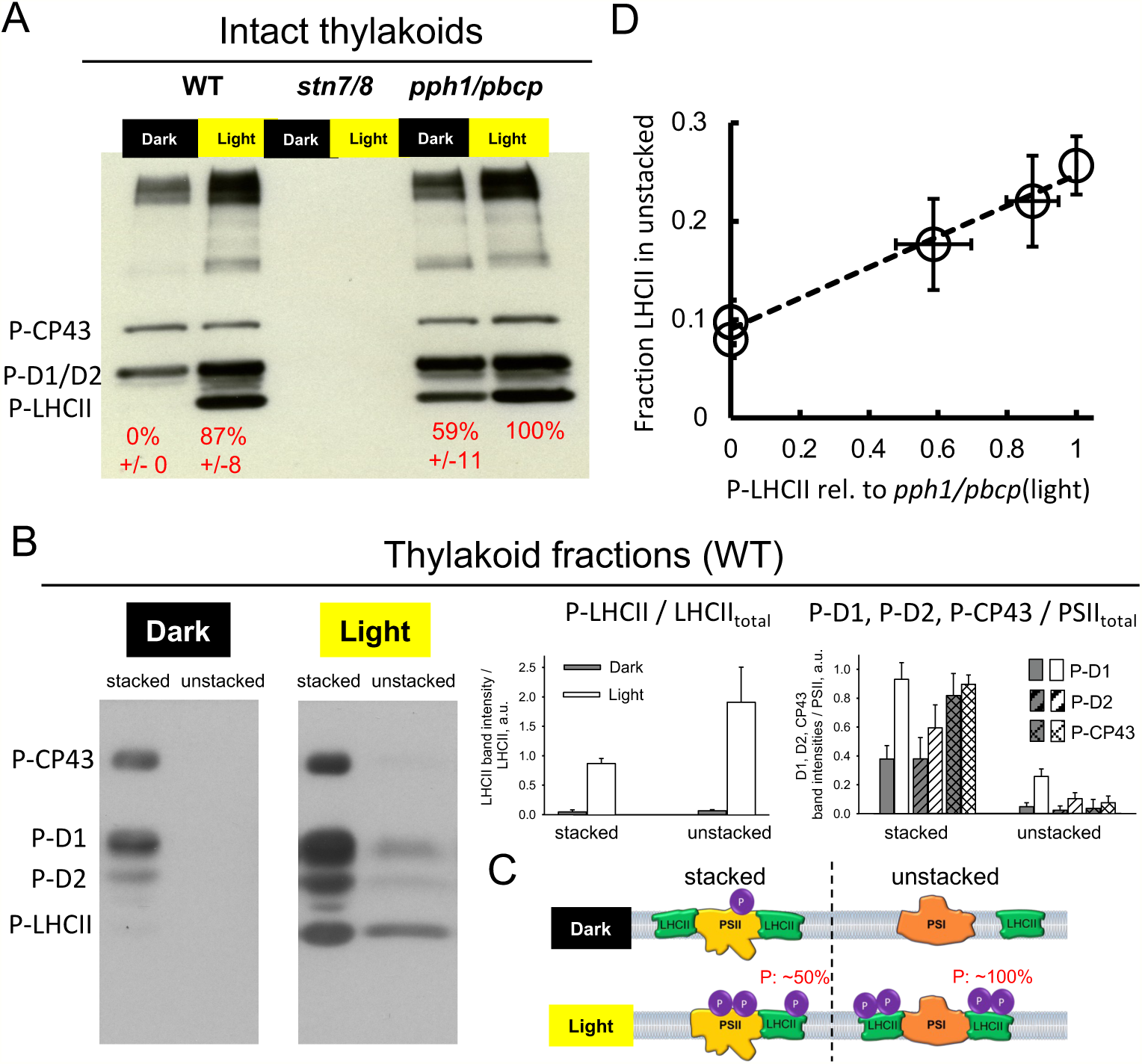
Protein phosphorylation levels in intact thylakoid membranes and thylakoid subdomains. **(A)** Representative Western immunoblot (WB) with phospho-threonine antibody with intact thylakoid membranes isolated from dark adapted or light-adapted (60 min, 120 μmol quanta m^-2^ s^-1^) wildtype and mutant plants. The identity of the phosphorylated protein complexes (P-CP43, P-D1/D2, P-LHCII) are indicated on the left. It is verified (Supplementary Fig.1) that the staining intensities are in a linear range. The red numbers at the bottom indicate the staining intensities of the P-LHCII band in percent relative to the *pph1/pbcp*-light sample that is expected to have full LHCII phosphorylation. Numbers represent the mean of two biological replicates with standard deviation. **(B)** left, examples for phospho-threonine WB for isolated stacked and unstacked thylakoid samples for dark and light. The histograms on the right give the WB band intensity normalized to the amount of LHCII (P-LHCII / LHCII_total_) or PSII (P-D1, P-D2, or P-CP43 / PSII_total_) subjected on the gel, i.e. it represents the phosphorylation level per protein. Mean with standard deviation from n = 3 biological repeats. **(C)** Cartoon comparing the phosphorylation levels of PSII and LHCII in stacked and unstacked thylakoid membranes for dark and light treated plants. **(D)** Linear relationship between the phosphorylation level of LHCII and the fraction of this complex localized in unstacked domains. The data shows the mean with standard deviation.

For the quantification of the protein phosphorylation levels in isolated stacked and unstacked WT thylakoid domains, the staining intensities of phospho-threonine antibody signals (Fig. 1B, left, for linearity of the signals see Supplementary Fig. 1) were normalized to the LHCII and PSII contents, respectively (Supplementary Fig. 2). This method allows for direct comparisons of the degree of phosphorylation per protein between stacked and unstacked domains (histograms in Fig. 1B). For LHCII, it turns out that switching to S2 causes an increase in LHCII phosphorylation in stacked thylakoids. In unstacked domains, however, this increase is even more pronounced, ending in a doubled phosphorylation level per LHCII compared to stacked regions. The PSII phosphoproteins D1, D2, and CP43 in stacked grana are already significantly phosphorylated under S1 that increases further under S2, mainly for D1 (Fig. 1B, right). In contrast to LHCII, the phosphorylation of PSII subunits under S2 in unstacked remains low. Fig. 1C summarizes the state transition induced protein phosphorylation changes. The doubled LHCII phosphorylation level in unstacked membranes compared to LHCII in stacked suggests that LHCII can escape stacked grana only if the protein is sufficiently phosphorylated, i.e. a certain threshold is exceeded.

### Are repulsive forces induced by phosphorylation sufficient for expelling LHCII out of stacked grana regions?

The observation that under S2 LHCII in unstacked is hyperphosphorylated raises the possibility that up to 12 additional negative charges from phosphate groups on LHCII (Supplemental text) generate sufficient electrostatic repulsion to force LHCII out of stacked grana by lateral diffusion. To test this possibility, a lateral force balance calculation was performed. To this end, the sum of repulsive electrostatic and attractive van der Waals forces were calculated between two ‘extra’ LHCII-trimers (Fig. 2A, see Supplemental text for details). The force balance analysis (Fig. 2) reveals that for unphosphorylated LHCII, van der Waals attraction outperforms electrostatic repulsion leading to net negative interaction energies (Fig. 2B and 2C, black stripes and blue line) and therefore LHCII aggregation. This is in line with recent experimental and theoretical data (Puthiyaveetil et al. 2017, Manna et al. 2023). The interaction energy of ∼ -3.3 k_B_T (Fig. 2D, energy expressed as thermal energy equivalents k_B_T; k_B_, Boltzmann constant; T, temperature in Kelvin) is in good agreement with experimental results of ∼ 5 k_B_T (Manna et al. 2023). These absolute numbers are relatively small (not far from thermal energy k_B_T) indicating no strong driving forces for LHCII aggregation. For 100% LHCII phosphorylation, the sum of forces is still in favor for LHCII aggregation, i.e. the van der Waals attraction is higher than the electrostatic repulsion. The minimum energy interaction level (*E_min_*) can be interpreted as the driving force for aggregation and the energy maximum (*E_max_*) the force for LHCII-LHCII separation. The population of the aggregated state (*P_agg_*) versus the population of the separated state (*P_sep_*) is given by the Boltzmann distribution:

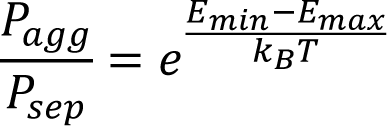

**Fig. 2.**
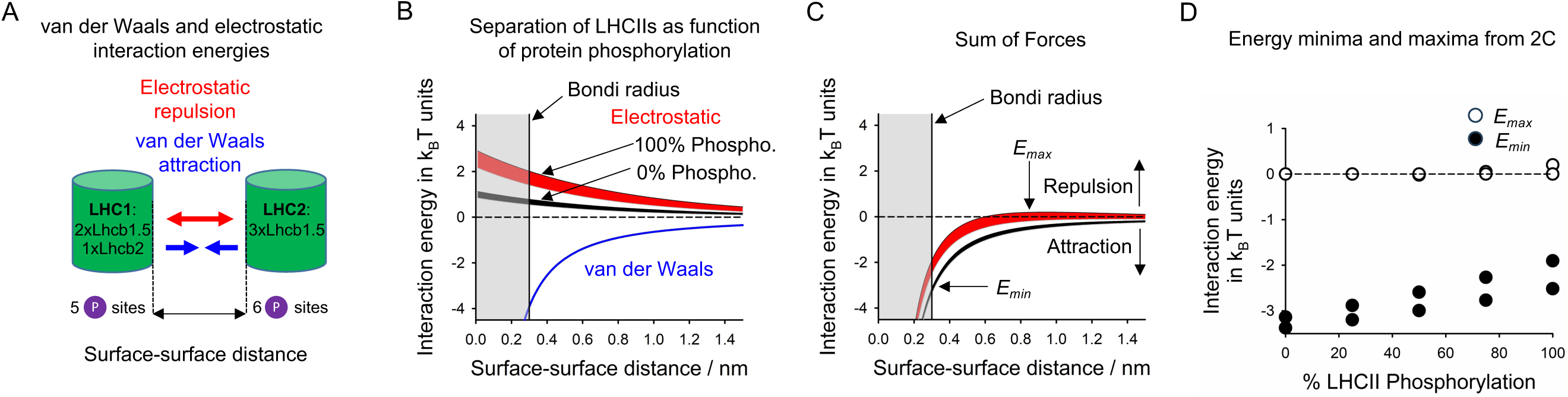
Force balance analysis for a pair of ‘extra’ LHCIIs. **(A)** Cartoon summarizing the subunit composition and phosphorylation sides for the two forms of ‘extra’ LHCIIs interacting by repulsive and attractive forces. **(B)** Calculated distance dependencies of electrostatic and van der Waals forces. Electrostatic repulsions are shown for the two extreme LHCII phosphorylation states (0% and 100%). The upper and lower boundaries of the black and red stripes represent 100 mM and 150 mM KCl concentrations in the stroma, respectively. The energies are represented in thermal energy units (k_B_T). Negative interaction energies correspond to LHCII-LHCII attraction positive values correspond to LHCII separation. The grey vertical stripe represents the physical hard-shell boundary of LHCIIs (Bondi radius), i.e. interaction energies must be counted from this distance on. **(C)** Sum of forces calculated from the curves in B. The maximal (*E_max_*) and minimal (*E_min_*) energy levels are indicated. These energy levels represent the energy for aggregation (*E_min_*) or separation (*E_max_*). **(D)** Summary of maximal and minimal energy values as a function of different LHCII phosphorylation levels in %. The values for 0% and 100% are derived from C. Energy curves for other phosphorylation levels are not shown. The pairs for values for each phosphorylation level is based on the two KCl concentrations (see stripes in C). For further details please see in the main text and in Supplemental text.

From the values in Fig. 2D it follows that the ratio of *P_agg_/P_sep_*changes from ∼25 at 0% phosphorylation to ∼10 at 100% phosphorylation. It can be concluded that the maximal LHCII phosphorylation does not generate enough driving force to initiate lateral migration of this protein complex from stacked to unstacked membrane domains. This conclusion is further strengthened by the fact that the enthalpy of aggregation increases with the number of LHCII (Manna et al. 2023), i.e. a situation realized in native grana. An increased number of LHCIIs will also lower the entropy for LHCII-LHCII interaction (Manna et al. 2023). The entropy counteracts LHCII aggregation. Finally, LHCII aggregation is energetically further stimulated by a pH drop (Manna et al. 2023) that is expected in illuminated plants as the lumen acidifies. All these observations indicate that the lack of driving force for a separation between phospho-LHCIIs seen in Fig. 2 is also realized in native membranes. If LHCII phosphorylation is not triggering LHCII lateral mobilization in grana under S2, then the question arises how phospho-LHCII can escape the protein network in stacked thylakoid membranes. The most likely explanation is that partial vertical unstacking of grana stacks under S2 (Kyle et al. 1984, Wood et al. 2018, Gatry et al. 2023) releases P-LHCII complexes that can then interact with PSI forming the LHCII-LHCI-PSI state transition complex. This is analyzed in the next section.

### Impact of protein phosphorylation on photosystem II and photosystem I supercomplex stability and formation

Under S2 conditions, part of the PSI-LHCI complexes can functionally attach to LHCII forming a PSI-LHCI-LHCII ‘state-transition complex’ (Pesaresi 2009, Suorsa 2015, Benson 2015). Furthermore, in stacked grana of dark-adapted Arabidopsis plants, most of the PSII is organized as a C2S2M2 supercomplex (C2, PSII core dimer; S, strongly bound LHCII; M, moderately bound LHCII) (Fig. 3A, Caffarri 2009, Kouril 2013, Koochak 2019). Evidence exists that reversible protein phosphorylation controls the assembly of both PSII and PSI supercomplexes (Leoni 2013, Dietzel 2011, Pan 2018) although this view was challenged for the PSII supercomplexes (Wientjes 2013). The PSII supercomplex quantifications based on BN PAGE combined with dot blot analysis (Koochak 2019) shows that under S1 to S2 transition ∼15% of C2S2M2 supercomplexes dissembles into smaller C2S2 and C2 in stacked domains (Fig. 3A). However, a same level of partial disassembly of C2S2M2 is also visible in both kinase and phosphate mutants, respectively suggesting that the partial PSII supercomplex disassembly is not phosphorylation but light-dependent (Suorsa 2015). In contrast to stacked thylakoid domains, ∼85% of PSII in unstacked domains is organized as a monomeric core (C) or CP43-free core (CP43-C) (Fig. 3B, Koochak 2019, Bassi 1995). Illumination leads to a partial conversion of CP43-C to C that could be a sign for an activated PSII repair cycle that requires thylakoid-generated ATP by (low) light.

**Fig. 3.**
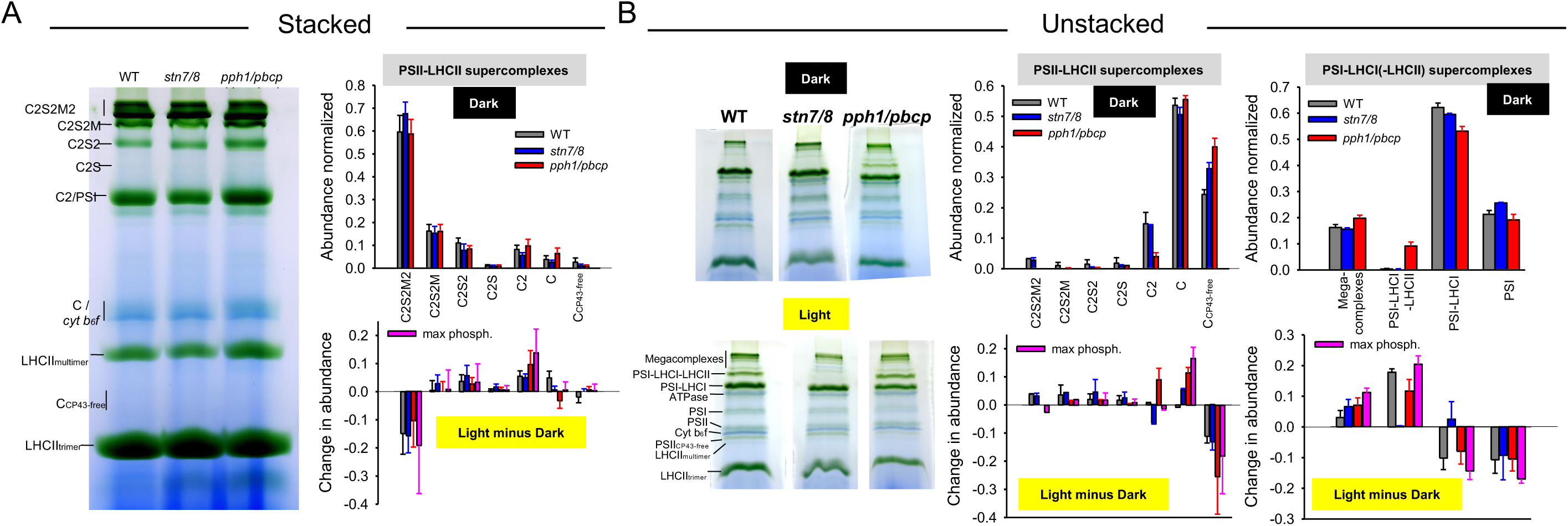
Photosystem supercomplex quantifications in stacked and unstacked thylakoid membranes. **(A)** Quantification of PSII supercomplexes for stacked grana. Left, representative blue native PAGE gel with labeled bands. The histogram to the top shows the fraction of each PSII complex determined from dot blot WB analysis of the cut-out gel bands with a highly specific D1 antibody as described in detail in (Koochak 2019). The histogram gives specific light-induced changes is PSII complex abundances. The data represents the mean with standard deviation from n = 3 to 5 biological repeats. No statistically significant changes between the three genotypes (student’s t-test). **(B)** Quantification of PSI and PSII supercomplexes for unstacked thylakoid membranes. Left, representative examples of BN-PAGE gels for dark and light adapted samples. The identity of the bands is indicated for the WT-light gel. The histogram in the middle shows PSII complex abundances derived in the same way as in (A). The histogram to the right shows the quantification of the PSI mega- and supercomplexes determined directly from the gel as described in the Method section. The data shows the mean with standard deviation from n = 3 biological repeats. The maximal phosphorylation induced change (pink bars) is the difference between *pph1/pbcp*-light minus *stn7/8*-dark since for these samples show maximal and no LHCII phosphorylation (see Fig. 1).

From a quantitative analysis of BN PAGE gels of unstacked thylakoid membranes (Fig. 3B, far right), it follows that in dark-adapted WT as well as in *stn7/8* plants with no LHCII phosphorylation (Fig. 1), ∼17% of PSI are assembled in (PSI-PSII-LHCI-LHCII) megacomplexes, ∼60% in PSI-LHCI complexes, and ∼20% in PSI. In dark adapted *pph1/pbcp* plants with ∼60% LHCII phosphorylation (Fig. 1), the abundance of the PSI-LHCI-LHCII ‘state-transition complex increased by ∼10%. In WT under S2, an additional ∼20% of PSI binds LHCII increasing the PSI-LHCI-LHCII/megacomplex abundance from 17% in S1 to 37% (Fig. 3 far right bottom) in agreement with previous reports (Benson 2015). Comparing light-induced PSI complex dynamics in WT with the two mutants reveals that the formation of PSI-LHCI-LHCII/megacomplexes is phosphorylation dependent (Fig. 3B, far right bottom), i.e. it is missing in *stn7/8* (no phosphorylation) and stronger in *pph1/pbcp* (hyperphosphorylation). The maximal phosphorylation-triggered change was determined by subtraction data for the *pph1/pbcp*-light samples from *stn7/8*-dark (see phosphorylation levels in Fig.1). It appears that maximal LHCII phosphorylation converts ∼1/3 of PSI/PSI-LHCI into PSI-LHCI-LHCII/megacomplexes in unstacked thylakoid domains (pink bars in Fig. 3B far right bottom) leading to half of PSI organized in LHCII-containing complexes. Overall, the results suggest that protein phosphorylation has no direct impact on PSII supercomplex stability in stacked grana but is required for the binding of LHCII to PSI-LHCI complexes in unstacked domains (Pesaresi 2009, Suorsa 2015, Wientjes 2013).

### Protein and chlorophyll redistribution between stacked and unstacked thylakoid membranes induced by state transition

The stacking of part of the thylakoid membranes to grana is accompanied by pronounced heterogeneity in protein complex distribution. The protein distribution analysis in Fig. 4A and Supplementary Fig. 3 shows an enrichment of LHCII and PSII in stacked domains whereas the abundance of PSI and ATPase is higher in unstacked thylakoid membranes (Staehelin 1996, Albertsson 2001, Koochak 2019, Pribil 2014). Within the error margin of the experiment, transition from S1 to S2 leads to no changes in the lateral distribution for PSII, PSI, and ATPase (dashed bars in Fig. 4A and Supplementary Fig. 3) but to a statically significant redistribution of 12% of LHCII from stacked to unstacked thylakoid membranes in WT. The lateral redistribution of LHCII in S2 from stacked to unstacked domains is in line with older reports (Kyle et al. 1983, Allen 1992) but in contrast to more recent publications where no such LHCII reorganization was seen (Wood et al. 2018). Inspection of the LHCII phosphorylation level reveals that in Wood et al. 2018, LHCII is already significantly phosphorylated in S1 in contrast to complete lack of LHCII phosphorylation in our study (Fig. 1). A likely explanation for the different phosphorylation levels in S1 and as a consequence thereof a different LHCII reorganization behavior, are different dark treatments in both studies. In our study, plants for S1 were dark adapted overnight whereas Wood et al. 2018 used a shorter dark-incubation time of 1 hour.

**Fig. 4.**
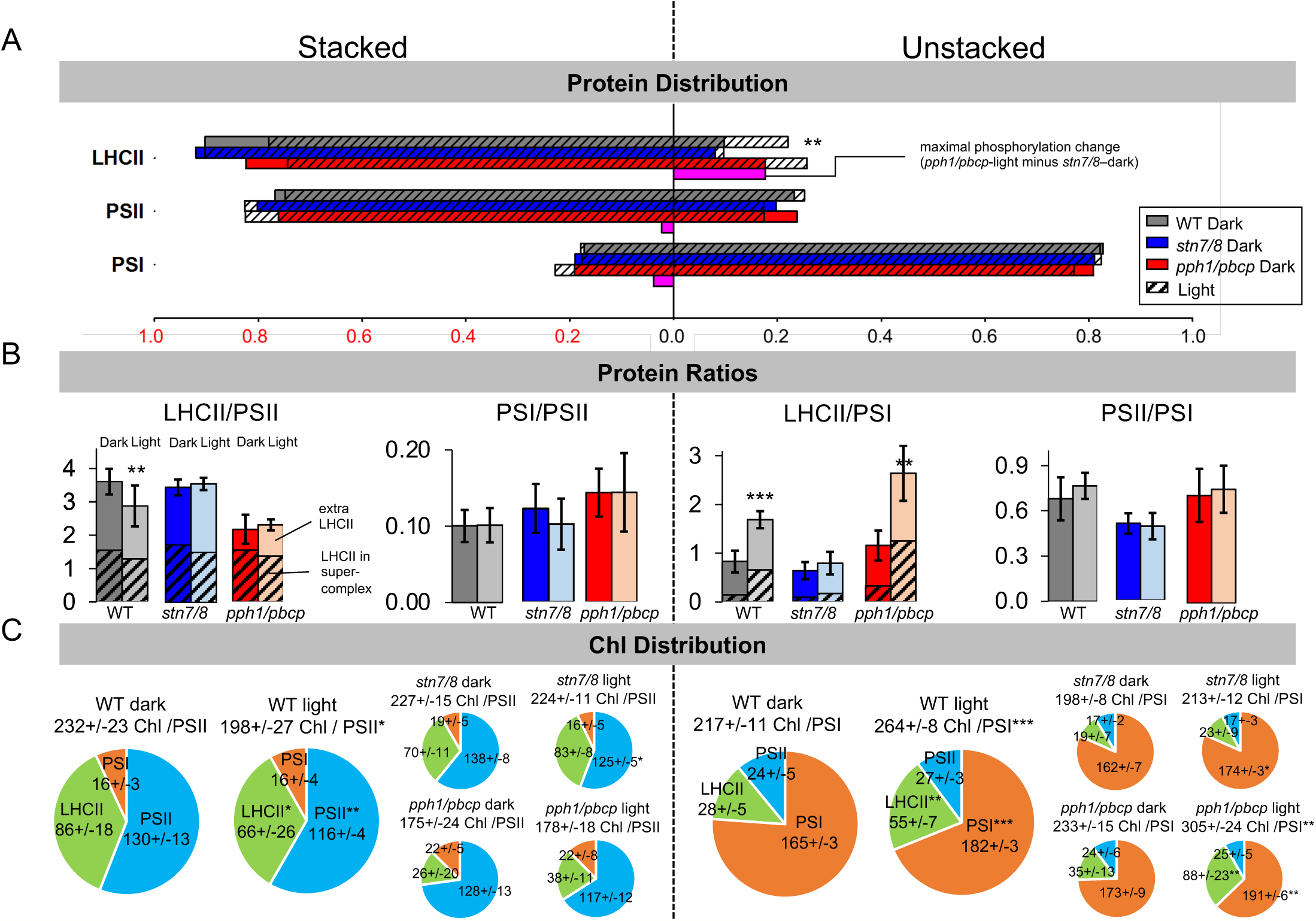
Protein and Chl distribution dynamics between stacked and unstacked thylakoid subdomains. **(A)** Lateral distribution of LHCII, PSII, and PSI between stacked and unstacked regions for the three genotypes. Solid bars give distributions for dark-adapted while shaded bars for light-adapted samples. Note that only for LHCII, a statistically significant redistribution induced by a S1 to S2 transition is apparent. For ATP synthase data, see Supplementary Fig. 3. The data shows the mean values from n = 3 to 6 (LHCII), n = 3 to 11 (PSII), and n = 3 to 8 (PSI) biological repeats. For clarity, error bars are omitted. Standard deviations are given in Table S1. **(B)** Protein complex ratios in stacked and unstacked domains for S1 and S2. The shaded bars show the fraction of LHCII organized in supercomplexes with PSII and PSI, respectively. ‘Extra LHCII’ indicates the fraction of LHCII not bound in supercomplexes. The data shows the mean values with standard deviations from n = 3 to 9 (LHCII/PSII), n = 3 to 9 (PSI/PSII), n = 3 to 6 (LHCII/PSI), and n = 3 to 8 (PSII/PSI) biological repeats. **(C)** State transition-induced dynamics of Chl distribution between PSII, PSI, and LHCII in both thylakoid domains. The number of Chls per protein complex were calculated from Fig. 4B and Table S2. The data shows the mean with standard deviations from n = 3 to 5 biological repeats. Significance levels: *, p<0.05; **, p<0.01, ***, p< 0.001.

Examining the behavior of the kinase and phosphatase mutants indicates that the lateral LHCII relocation is phosphorylation dependent since *stn7/8* has the lowest and *pph1/pbcp*(light) the highest LHCII abundance in the unstacked thylakoid area. A linear relation between LHCII phosphorylation level and LHCII abundance in unstacked (Fig. 1D) provides strong evidence for this hypothesis. The maximal lateral redistribution capacity for the entire LHCII pool induced by state transition is 18% (pink bar in Fig. 3A) in accordance with the literature (Allen 1992, Staehelin 1996). The LHCII distribution behavior in Fig. 4A is confirmed by Chl a/b ratios (Supplementary Fig. 4). A consequence of lateral protein reorganizations is that the LHCII/PSII ratio in stacked grana in WT declines from 3.6 to 2.9 with a concomitant increase of the LHCII/PSI in unstacked from 0.8 to 1.7 (Fig. 4B). As seen for *pph1/pbcp* (light), the LHCII/PSII ratio can decrease maximally to 2.2 which is paralleled by an increase of the LHCII/PSI ratio to 2.6. As expected from the unchanged photosystem distributions in Fig. 4A, the PSII/PSI and PSI/PSII ratios in stacked and unstacked, respectively, do not change significantly.

Both photosystems show a different behavior for supercomplex formation (shaded bars in Fig. 4B) as a function of the total LHCII/photosystem ratios (non-shaded bars in Fig. 4B). The fraction of PSII-LHCII supercomplexes is fairly constant although the total LHCII/PSII ratios changed significantly for the three genotypes (Fig. 4B, left). In contrast, the histograms in Fig. 4B (right) suggest a strong correlation between PSI-LHCII mega-/supercomplexes formation and the total LHCII/PSI ratio. This correlation is visualized in Supplementary Fig. 5 indicating that both a full LHCII phosphorylation and an increased LHCII/PSI ratio is required for an efficient PSI-LHCII complex formation.

The protein complex quantifications shown in Fig. 4B enables detailed analysis of changes in Chl distribution induced by state transition (Fig. 4C). In WT, transition from S1 to S2 leads to a decrease in the total Chl/PSII ratio from 232 to 198 in stacked grana that is accompanied by an increase of Chl/PSI from 217 to 264 in unstacked thylakoid regions. A more specific examination of Chls for PSII in stacked and PSI in unstacked was done by subtracting Chls from PSI in stacked and of PSII in unstacked revealing that the number of Chl per PSII in WT decreases by 16% from 216 (86+130, S1) to 182 (S2). At the same time, the antenna size of PSI in unstacked increases by 23% from 193 (28+165, S1) to 237 (S2) in accordance with previous reports (Benson 2015, Lunde 2000, Kim 2015).

The higher unequal Chl/PSI increase compared the Chl/PSII decrease in S2 can be explained by the higher abundance of PSII relative to PSI. In detail, from the PSII/PSI ratio of 1.58 determined for the same plant material (Svoboda 2023) and the photosystem distribution in Fig. 4A, it follow a PSII_stacked_/ PSI_unstacked_ ratio of 1.46 (S1) and 1.44 (S2). These ratios are almost identical to the inverse of the S2 induced PSI/PSII antenna size changes of 1.43 (23% / 16%, see above). The similarities of these numbers indicate that 16% of Chls detaching from PSII in grana bind quantitatively to PSI unstacked regions leading to a larger 23% PSI antenna size increase because there are more PSII than PSI complexes in thylakoid membranes.

### Functional implications of lateral LHCII redistribution induced by state transition

The functional antenna sizes for PSII and PSI were analyzed by low temperature Chl fluorescence spectroscopy both with whole leaves and intact thylakoid membranes (Fig. 5). At 77K temperature, the emission spectra show two main peaks, one around 690 nm corresponding to LHCII-PSII and one around 730 nm corresponding to LHCI-PSI (Andreeva 2003). The 690 nm peak consists of contributions from the CP43 subunit of PSII (∼684 nm) and from CP47 (∼693 nm) whereas the 730 nm emission contains signals from LHCI bound to PSI (∼720 nm) and from the PSI core (∼733 nm) (Andreeva 2003). Mathematical fitting of the spectra with Gauss functions (Andreeva 2003) leads to the separation of emission spectra from LHCII-PSII (F684 + F692) and LHCI-PSI (F720 + F733) (Supplementary Fig. 6). Since the area under each fluorescence component is proportional to the light absorbed, the spectral fitting with Gauss functions provides direct measurements of the total antenna sizes of PSII and PSI, respectively. Results for intact leaves and isolated thylakoid membranes provide consistent results (Fig. 5A and 5B) indicating that both S1 and S2 states were preserved during thylakoid isolation.

**Fig. 5.**
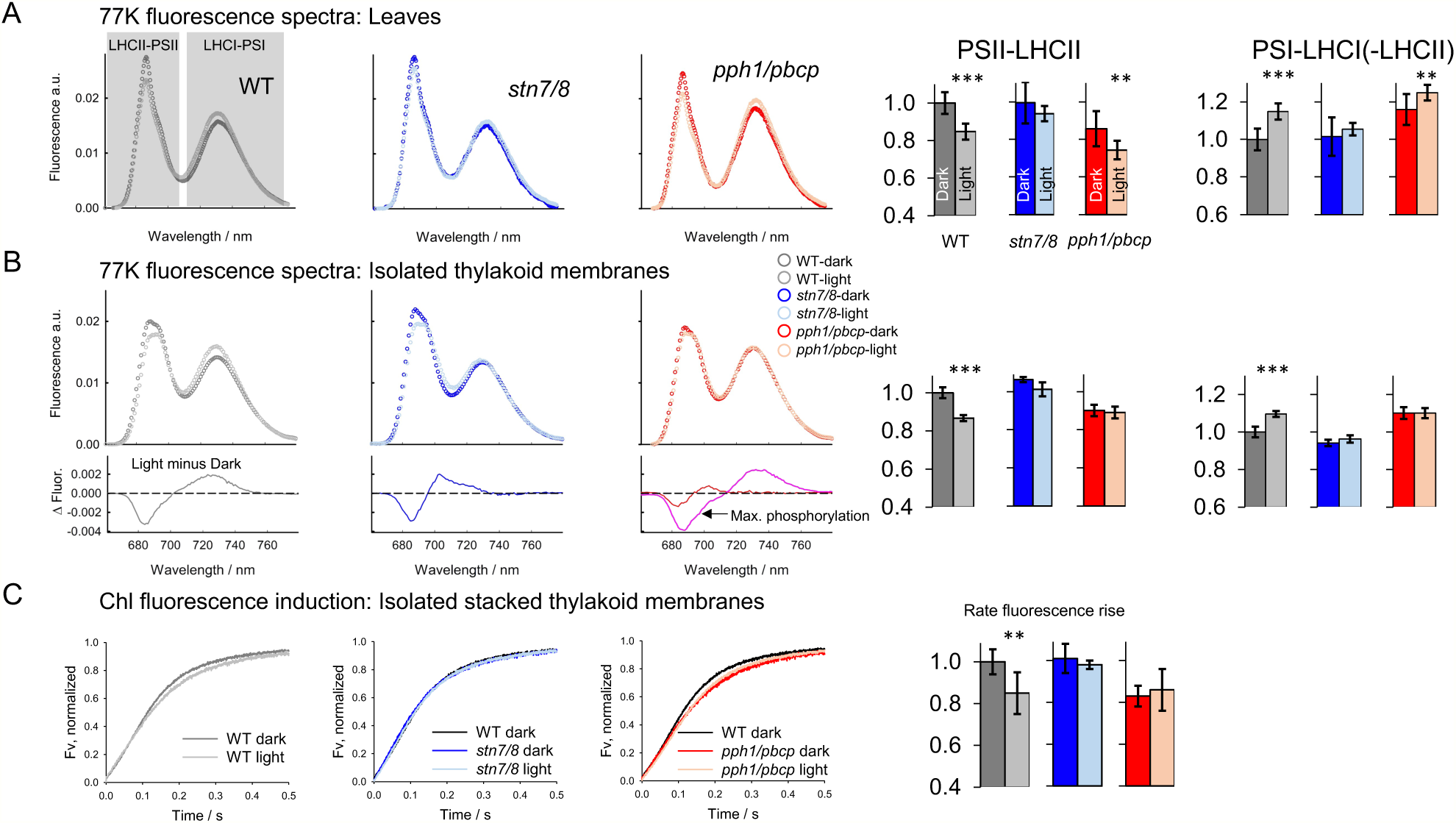
State transition triggered changes in light harvesting functions by PSII and PSI. Representative examples of low temperature Chl fluorescence spectra for leaves **(A)** and intact thylakoid membranes **(B)** for all three genotypes. Left, examples of fluorescence spectra. The spectra were mathematically fitted with four Gauss functions to separate LHCII-PSII (Gauss function for CP43 and CP47) and LHCI-PSI (Gauss functions for PSI core and PSI-LHCI) components (see Supplemental Fig. 8 for details). The area under the spectra were normalized to unity allowing the comparison between different samples. The areas under the fitted lines are proportional to the antenna cross sections of the individual photosystems. PSII-LHCII are the sum of the CP43 and CP47 areas, PSI-LHCI(-LHCII) are the sum of PSI-core and PSI-LHCI). These total PSII and PSI areas are given in the histograms to the right (normalized to WT-dark area). The light minus dark difference spectra shown at the bottom panel of **(B)** visualize the change in antenna cross sections, i.e. redistribution of excitation energy from LHCII-PSII (negative) to LHCI-PSI (positive). The maximal phosphorylation change (pink line) represents the difference *pph1/pbcp*-light minus *stn7/8*-dark. The data shows the mean values with standard deviations from n = 6 to 15 biological repeats for the leaves and n = 3 to 6 for isolated thylakoid membranes. **(C)** Representative examples of Chl fluorescence induction curves recorded at room temperature with isolated stacked grana preparations. Kinetics were measured in the presence of 60 μM dichloromethylurea (DCMU, blocks electron transport from QA to QB) and 1 mM diphenylcarbazide (PSII electron donor bypassing the oxygen evolving complex). The kinetics show the transition from Fo (time point 0.0 s) to the maximal fluorescence level Fm (normalized to 1). Fv = Fm – Fo. The speed of this Fv transition is proportional to the antenna cross section of PSII. The rate of Fv increase is expressed as the reciprocal time point when 60% of Fv is reached and summarized in the histogram to the right (normalized to WT-dark). The data shows the mean values with standard deviations from n = 3 to 7 biological repeats. Significance levels: *, p<0.05; **, p<0.01, ***, p< 0.001.

In WT, a S1 to S2 transition leads to a smaller LHCII-PSII antenna size and a corresponding increase in the (LHCII-)LHCI-PSI antenna (histograms in Fig. 5A and 5B). The redistribution of excitation energy from PSII to PSI is readily visible in the difference plot (Light minus Dark). Quantification of the area below the LHCII-PSII component reveals a ∼15% decrease in the PSII antenna size (left histograms in Figs. 4A and 4B) in accordance with the change in Chl/PSII ratio in stacked grana (∼16%, see above). A ∼15% decrease in the functional PSII antenna size induced by state transition for WT is also apparent from Chl fluorescence induction measurements at room temperature of isolated stacked grana (Fig. 5C). The rate of the fluorescence rise (histogram in Fig. 5C) derived from these curves is a measure of the PSII antenna size (Rappaport 2007). The congruence of antenna size changes examined by the two methods indicates that the changes in PSII antenna cross section is localized in stacked thylakoid membranes, as expected. Extension of the functional analyses to the two double mutants shows that the changes in antenna cross sections are dependent on LHCII-phosphorylation.

What are the implications of the Chl redistribution by state transition for the electron flux balance through both photosystems? Electron transport rates (ET) can be calculated from photochemical quantum yields (ϕ_PSII_, ϕ_PSI_), the proportion of incident light that is absorbed by the leaf (*A_leaf_*), the fraction of absorbed light received by PSII or PSI, respectively (*Fraction_PSII_*, *Fraction_PSI_*), and the photosynthetically active photon flux density (PPFD) by the equations (Baker 2008):

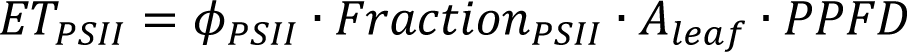

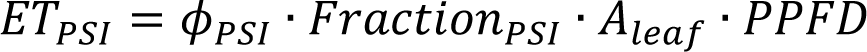

The fraction of absorbed light received by each photosystem is given by the photosystem ratio (*R_PSII_/R_PSI_*) and the number of chlorophylls bound per each reaction center (*Chl_PSII_, Chl_PSI_*):

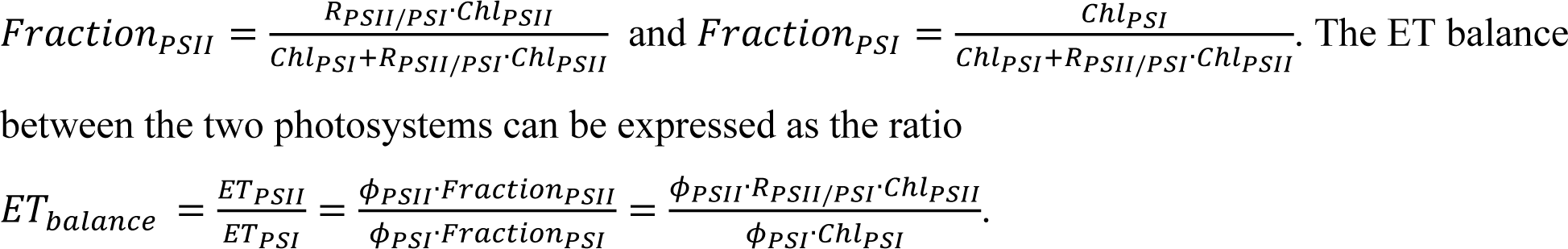

The photochemical yields for WT for both states are 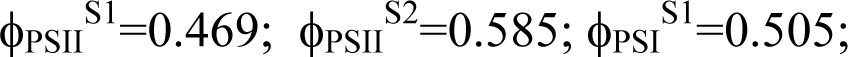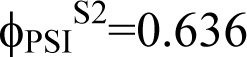 (Supplementary Fig. 7). It follows that the electron transport balances for S1 and S2 are (using Chl and *R_PSII_/R_PSI_*data from the previous section):

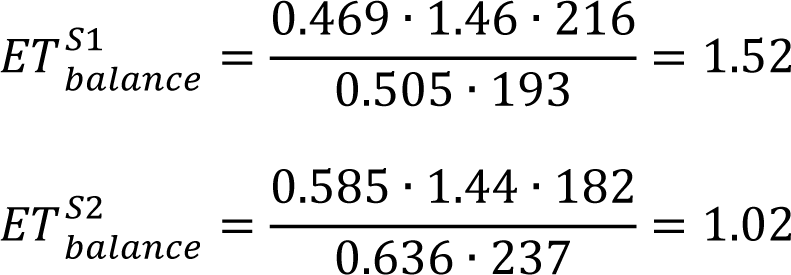

Thus, the transition from S1 to S2 leads to perfect balanced electron transport fluxes through PSII and PSI starting from a strongly PSII-favoring regime in S1 (52% higher relative ET rate). These numbers demonstrate the physiological significance of state transition for synchronizing electron transport rates through both photosystems under low light. This leads no room for additional cyclic electron transport around PSI in accordance with observations that the CET activity is low under low light (Morales 2018, Sagun 2019).

As expected from LHCII protein data, the Chl redistribution from PSII-stacked to PSI-unstacked is phosphorylation dependent: the Chl/PSII ratio is large (∼210) without LHCII phosphorylation (*stn7/8*) and small (∼155) under hyperphosphorylation (*pph1/pbcp*) with the opposite change for PSI in unstacked (∼180 for *stn7/8*-dark and ∼280 for *pph1/pbcp*-light). Consequently, the mutants cannot fine tune electron transport through both photosystems in the light with *stn7/8* showing ∼40% higher ET rate of PSII relative to PSI whereas the PSI rate is ∼20% higher in *pph1/pbcp* (Supplemental Fig. 7). The lack of fine-tuning ET fluxes through both photosystems is likely the reason for the severe fitness handicap of *stn7/8* and *pph1/pbcp* under field conditions (Kühlheim 2002, Frenkel 2007).

### A quantitative view on state transition in Arabidopsis

Figure 6 summarizes the maximal structural and functional changes induced by a S1 to S2 state transition by comparing the two extreme cases examined in this study: no LHCII phosphorylation in *stn7/8* (S1) and hyperphosphorylation in *pph1/pbcp*-light (S2). We choose these extremes to demonstrate the maximal potential of state transitions in higher plants. In S1, each PSII-dimer in stacked grana is associated with up to seven LHCII-trimers with half of them integrated in LHCII-PSII supercomplexes and the other half as ‘extra’ LHCII. In unstacked thylakoid domains, ∼85% of PSI is organized as PSI-LHCI or PSI w/o LHCI and 15% as LHCII-containing super-/megacomplexes. In the extreme S1, the PSII/PSI antenna size ratio is 1.68 (calculated from bottom table Supplemental Fig. 7) favoring a strongly biased PSII ET over PSI (previous section). This will lead to an over-reduction of the PQ-pool with potential photodamage (Grieco 2012). A shift to S2 causes a doubled phosphorylation level of LHCII in unstacked compared to their counterparts in stacked area, indicating that full LHCII-phosphorylation is required for LHCII to escape from grana (see also Fig. 1).

**Fig. 6.**
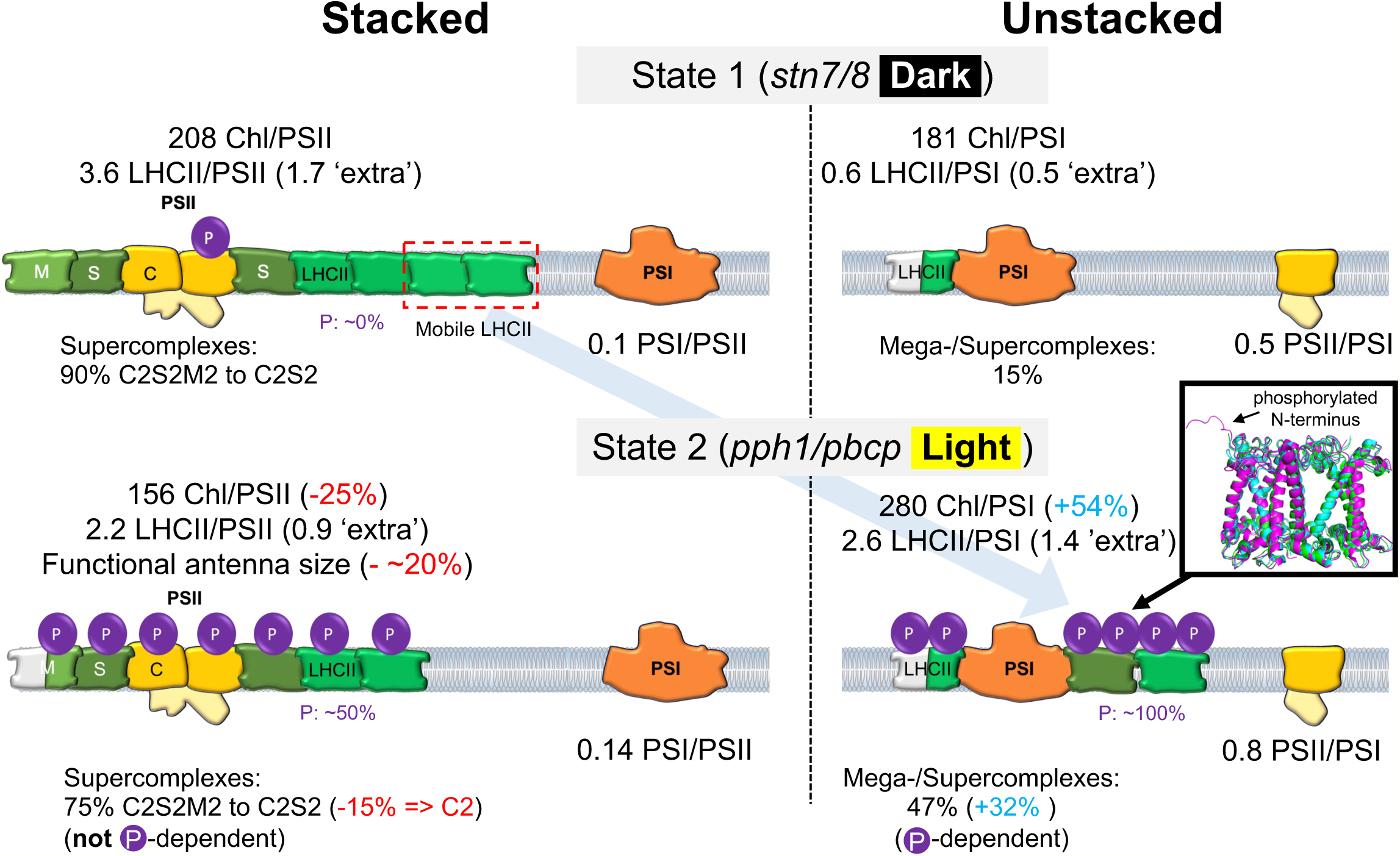
Summary of the structural and functional status of the light harvesting apparatus in stacked and unstacked thylakoid domains. The survey represents extreme cases for S1 as seen in *stn7/8*-dark (no LHCII-phosphorylation) and S2 as seen in *pph1/pbcp*-light (maximal LHCII-phosphorylation). The figure shows quantifications of the #Chls per photosystem, ratio of LHCII to photosystem, #LHCII organized in supercomplexes and ‘extra’ LHCII, functional antenna cross sections, and LHCII phosphorylation levels. The inset compares structures of non-phosphorylated LHCII-trimers organized in the C2S2M2 in stacked grana (S-trimer in green, pdb# 5XNL; M-trimer in cyan, pdb# 5XNL) or of phosphorylated LHCII in the PSI-LHCI-LHCII ‘state transition complex’ in unstacked thylakoid membranes (pdb# 5ZJI). The structural comparison indicates that LHCII-phosphorylation and attachment to PSI leads to distinctive changes in the N-terminus adopting a structured conformation (highlighted by the black arrow) whereas the structure of the rest of the protein remains largely unchanged.

In the extreme S2, up to two ‘extra’ LHCII (or ∼20% of total LHCII) redistribute from stacked areas decreasing the antenna size of PSII by ∼25% paralleled by a ∼50% increased PSI antenna size. As described above this redistribution of hyperphosphorylated LHCII from PSII to PSI is probably not caused by lateral migration of the protein but by vertical grana unstacking. However, on the lateral in-membrane level, phospho-LHCII still has to unbind from PSII supercomplexes and bind to PSI leading to form LHCII-PSI mega-/supercomplexes (∼1/3 increase). Since an increase in the overall electrostatic repulsion is likely not generating sufficient power for phospho-LHCII to unbind from other LHCIIs and LHCII-PSII supercomplexes (see above and Mao et al. 2023), the question is what is the mechanism for PSII-unbinding and concomitant PSI-binding in S2? On the molecular level, the only significant structural change in LHCII triggered by its phosphorylation is apparent for the N-terminus (Fig. 6, inset, Supplementary Fig. 8). The rest of the protein structure remains virtually unchanged. Therefore, a viable scenario is that conformation changes of the phospho-LHCII N-terminus lower the binding affinity to LHCII-PSII in grana allowing phospho-LHCII to explore and establish new interactions with PSI in unstacked thylakoid domains. This scenario would favor a ‘molecular recognition’ concept (Allen 2019, Wood 2020). Whether this shift of binding affinities of phospho-LHCII between PSII and PSI in S2 is realized in the grana margins postulated as potential mixing zone for both photosystems (Rantala et al. 2020), has to be verified in future studies.

## Methods

### Plant growth conditions

6-to 8-week-old *Arabidopsis* (Col-0) and mutants (*stn7/8* and *pph1/pbcp*) were grown in soil in a growth chamber at 21°C and 60% relative humidity with 150 μmol quanta m^-2^ s^-1^ light of 9 h/15 h (day/night) photoperiod.

### Intact thylakoid, stacked and unstacked thylakoid extraction

A minimum of 4 plants were combined for intact thylakoid isolation. Plants were dark-adapted overnight and were treated with 120 μmol quanta m^-2^ s^-1^ light for 1 hour to induce state transition before thylakoid isolation. Leaves were homogenized with a blender in a grinding buffer that contained final concentrations of 20 mM Tricine (pH 8.4), 0.4 M Sorbitol, 10 mM EDTA, 10 mM NaHCO_3_, and 0.15% (w/v) of BSA. The homogenate was filtered through 1 layer of miracloth and 8 layers of cheese cloth to remove leaf debris. The filtrate was centrifuged at 2000 x g for 2 minutes. The supernatant was discarded and the pellet was resuspended in a large volume (∼40 mL) of chloroplast lysis buffer (25 mM HEPES (pH 7.5), 40 mM KCl, and 7 mM MgCl_2_). The resuspended material was incubated on ice for 10 minutes, followed by centrifugation at 2000 x g. The thylakoid pellet was washed once by resuspension in storage buffer (50 mM HEPES (pH 7.5), 0.1 M Sorbitol, 15 mM NaCl, and 10 mM MgCl_2_) and by subsequent centrifugation at 1485 x g. The pellet was finally resuspended in storage buffer and kept on ice. Freshly prepared thylakoids were used for a digitonin-based thylakoid fractionation. Isolated thylakoid membranes (0.6 mg/ml) were mixed with an equal volume of ice-cold 1% (w/v) digitonin solution and incubated at RT in dark for 10 minutes. The unsolubilized thylakoids which were pelleted after centrifugation at 1000 x g for 1 minute were discarded. The supernatant was subsequently centrifuged at 40000 x g for 30 minutes. The stacked membrane was retrieved as a pellet and was resuspended in a small volume of storage buffer. The supernatant consisted of unstacked thylakoid regions. All the centrifugation steps were at 4°C. The detailed method was published in (Koochak et al. 2019).

### SDS-PAGE Gel Electrophoresis analysis

LHCII: LHCII proteins in samples were determined on Coomassie stained SDS-PAGE gels. For quantification, isolated protein standards were compared with the samples after running on the same gel. The densitometric analysis of LHCII protein bands on the gels were performed using the Image-Pro Plus software and the concentrations of LHCII proteins were determined as mmol/mol Chl. The detailed method is described in (Svoboda et. al, 2023).

ATPase: The isolated ATPase protein standards (very kind gift from Dr. Georg Groth, Heinrich Heine University of Düsseldorf, Germany) dilution series were run on the same gel with the unknown sample on a Coomassie-stained SDS-PAGE gel (16% Tris-Glycine). To determine ATPase protein content, densitometric analysis of the staining of the ATPase β-subunit on the gel was performed. Using Image-Pro Plus software, the amount of the unknown sample (in mol) was determined from a regression curve generated from the dilution series of the standard ATPase β-subunit. ATPase concentration was then presented as mmol ATPase/mol Chl (Svoboda et. al, 2023).

### Protein phosphorylation

The phosphorylation status of PSII core and LHCII proteins were measured by Western blot analysis applying a phospho-threonine-specific antibody. Thylakoids and thylakoid fractions were resolved in 11% SDS-PAGE based on Laemmli. Dilution series of thylakoid membranes were used in each gel to check for signal linearity. Proteins were transferred on PVDF membrane in Towbin buffer without methanol in a Mini Trans-Blot cell (350mA/60min). Membranes were blocked with 5% BSA in TBST buffer for 60 min at room temperature and washed three times with TBST. Incubation with polyclonal primary Anti Pospho-Threonine antibody (Cell Signaling) diluted to 1:1000 in TBST was done overnight at 4°C. Membranes were washed with TBST and incubated with secondary antibody (Amersham ECL Rabbit IgC, HRP-linked) diluted to 1:50,000 in TBST at room temperature for two hours and washed with TBST. Amersham ECL Western Blotting Detection Reagents in combination with Amersham chemiluminescence Hyperfilm ECL were used for protein detection. Hyperfilm sheets were developed in GBX (Carestream Dental) developer and fixer, dried and documented. Image analysis was done in ImagePro software.

### PSII supercomplexes quantification

The quantification was based on BN-PAGE combined with dot blot analysis. Thylakoid, stacked and unstacked, membranes (50 ug Chl) contained 1% α-dodecylmaltoside (α-DM) and 0.5% Coomassie blue-G were ran in 12% gradient acrylamide gels at 50 V, 350 A for 18 to 20 h. The protein bands containing PSII supercomplexes were cut out and eluted by a Model 422 Electro-Eluter (Bio-Rad, Hercules, CA, USA). The relative amount of PSII supercomplexes were quantified by dot immunoblot analysis with D1 antibody (Agrisera, AS05 084). The dot intensities were quantified by densitometric analysis using Image-Pro Plus software. The detailed method was published (Koochak et al. 2019).

### PSII quantification

Stacked and unstacked membranes were resuspended in BBY buffer (pH 6.5, containing 15 mM MES, 10 mM KCl, and 0.5 mM EDTA) and the absorption at a spectral range of 540 to 575 nm was recorded by a Hitachi U3900 spectrometer. The chlorophyll concentration in the cuvette was adjusted to 25 to 40 μM as determined by the maximum peak at 677.5 nm (spectral range: 600– 750 nm). Cyt *b559*, PSII was quantified as described in Kirchhoff et al. 2002 with minor modifications. The redox change was induced by incubating samples consecutively with 1 mM potassium ferricyanide for 1 min, 4 mM sodium ascorbate for 5 min, and 5 mM sodium dithionite for 5 min (Svoboda et. al., 2023).

### P700 measurements

Photosystem I levels in thylakoids and thylakoid fractions were determined spectroscopically by P700 difference absorption measurements at 705 nm. Samples were diluted to 30 µg of chlorophyll/ml in buffer containing: 10 mM HEPES, 5 mM MgCl_2_, pH 7.5. Sodium ascorbate, and methylviologen were added before measurement to a final concentration of 20 mM and 100 µM respectively. Intact thylakoids were briefly sonicated before measurement. Sodium ascorbate reduced P700 was completely oxidated with 100 ms long saturating light pulse and recorded. For further details, see Kirchhoff et al. 2002.

### Total chlorophyll quantification

The amount of total chlorophyll per PSII monomer (measured as *cyt b559*) was calculated by the equation:

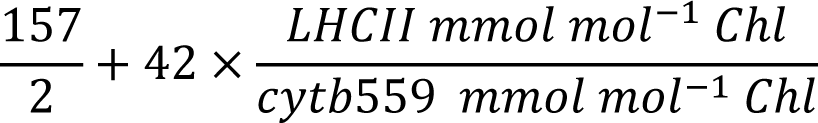

where 157 is the number of chlorophylls in a PSII dimer and 42 is the number of chlorophylls in LHCII (Cao et al., 2018).

### Low temperature (77 K) chlorophyll fluorescence spectroscopy

Leaf samples were frozen in liquid nitrogen, ground in a mortar to a powder, and then ground in the presence of ice-cold grinding buffer. The homogenate was filtered through a cell strainer (70 µm, Falcon) to remove leaf debris. The leaf filtrate or isolated thylakoid membranes were diluted in storage buffer (pH 7.5, pretreated with 22 units mL^-1^ glucose oxidase, 800 units mL^-1^catalase, and 4 mM glucose) to a Chl concentration of 10 μg/mL, and transferred into a glass tube. The glass tube with the leaf sample was shock-frozen in liquid nitrogen. The glass tubes were then transferred to a Dewar (Horiba Jobin Yvon) filled and pre-cooled with liquid nitrogen for measurement. Low temperature (77 K) fluorescence emission spectra were obtained with a Horiba Jobin Yvon FluoroMax 4 spectrofluorometer. The signal was recorded from 650 nm to 800 nm and averaged with 6 scans. Excitation wavelength was 435 nm. The 77 K spectra were fitted with four Gaussian curves to separate PSII (CP43 and CP47) and PSI (LHCI-PSII, PSI core) contributions (Andreeva et al. 2003). A global mathematical fitting routine includes simultaneous fitting of all WT and mutant 77K spectra. This was done separately for leaf samples and thylakoid membranes (see Supplemental Fig. 6). The areas under the PSII and PSI signals were determined providing a measure of the antenna sizes of both photosystems. All calculations were done by using SigmaPlot v11.0 software.

### Photochemical quantum yield of PSII and PSI

Measurements were performed using a MultispeQ fluorometer (PhotosynQ Inc.). Leaves attached to six-week-old plants were pre-illuminated for 20 sec (300 µE white light) to activate the PSI acceptor side (minimize acceptor side limitation), then exposed to 100 µE actinic light until steady-state was obtained (Supplemental Fig. 7). The steady-state fluorescence level (Fs) was measured with an amber 590 nm light and maximal fluorescence (Fm′) was measured by application of MT pulse (500ms) during steady-state. Photochemical quantum yield of PSII (ΦII) was calculated as (Fm′ – Fs)/Fm′ (Baker 2008). The photochemical quantum yield of PSI (ΦI) was derived from P700 signals measured as absorption change at 820 nm using a similar light protocol described above. After steady-state was obtained (Supplemental Fig. 7), P and Pm’ were measured by application of MT pulse (500ms), where P is P700^+^ level in steady state and Pm′ is P700^+^ maximal level in the MT pulse. Then, after 3 sec of 50 µmol quanta m^-2^ s^-1^ far-red light (FR) illumination, another MT pulse (500ms) was applied to measure the maximal P700 oxidation level Pmax). The minimal P700 oxidation level (Po) was determined by a 0.5 s FR interval followed directly after the MT pulse (Supplemental Fig. 7). ΦI was calculated as (Pm′ − P)/(Pm - P0) (Schreiber and Klughammer 2016).

### Chlorophyll fluorescence induction

Isolated stacked grana were used for Chl fluorescence induction and measured with a home-built fluorometer (Kirchhoff et al., 2004) at room temperature in the dark. 7 μg of Chl of stacked grana were mixed with 1.3 ml of BBY buffer (pH 6.5, containing 1 mM Diphenyl carbazide, DPC, Tang and Satoh 1985) in a square photometer cuvette. The sample was dark-adapted for 13 min and then 3-(3,4-dichlorophenyl)-1,1-dimethylurea (DCMU, 60 μM) was added in complete darkness. After the sample was incubated with DCMU for 30 sec, chlorophyll fluorescence variance (Fv) was measured with a green 530 nm light. The Fv was normalized to one. Fluorescence induction curves were analyzed with SigmaPlot 11.0 software. The minimum Chl fluorescence (Fo) was confirmed by adding potassium ferricyanide (FeCy, 3 mM) to the sample. (Koochak et al. 2019).

## Acknowledgments

We thank Dr. Mei Li (Chinese Academy of Sciences) for her very helpful discussions about phosphorylation-dependent structural changes of LHCII and preparing the inset in Fig. 6 and Supplementary Fig. 8. Furthermore, we like to thank Dr. Frank Müh (University of Linz, Austria) and Dr. Doran Raccah (University of Texas, Austin, USA) for their help to calculate lateral force balances between LHCIIs. This work was supported by a grant from the U.S. National Science Foundation MCB-BSF-1953570 (H.K.), MCB-BSF-1616982 (H.K.), and the US Department of Agriculture (ARC grant WNP00775).

## Author contributions

Conceptualization: HKi

Methodology: HMOO, HKo, VS, MK, HKi

Investigation: HMOO, HKo, VS, MK, HKi

Visualization: HMOO, HKo, MK, HKi

Funding acquisition: HKi

Project administration: HKi

Supervision: HKi

Writing – original draft: HKi

Writing – review & editing: HMOO, HKo, VS, MK, HKi

